# Structural basis for aphidicolin inhibition of human processive DNA synthesis

**DOI:** 10.64898/2026.10.19.760038

**Authors:** Ammar U. Danazumi, Muhammad Tehseen, Lingyun Zhao, Saleh Alshehri, Vlad-Stefan Raducanu, Samir M. Hamdan, Alfredo De Biasio

**Author notes:** These authors contributed equally (Ammar U. Danazumi, Muhammad Tehseen).

## Abstract

Eukaryotic DNA replication requires RNA–DNA primer synthesis by Pol α and processive lagging- and leading-strand synthesis by Pol δ and Pol ε, respectively, tethered to the PCNA clamp. Aphidicolin, a fungal diterpenoid, is a selective inhibitor of these B-family polymerases and a tool for inducing controlled replication stress, yet its structural mechanism has been defined only for the catalytic core of Pol α. Here we report cryo-EM structures of human Pol δ and Pol ε holoenzymes bound to primer-template DNA, PCNA and aphidicolin at 3.0 Å and 3.1 Å, respectively. In both, aphidicolin occupies the incoming dNTP pocket and flips the templating base outward, locking the fingers open and arresting synthesis while preserving the holoenzyme architecture. Aphidicolin occupies an architecturally conserved pocket in Pol α, Pol δ and Pol ε, yet Pol δ is approximately 4-5-fold more sensitive than Pol ε in reconstituted PCNA-dependent assays. We attribute this to a grip beyond the templating base that anchors the template strand in Pol δ but engages it only loosely in Pol ε, allowing the strand to escape the aphidicolin trap. Selectivity therefore resides not in the inhibitor pocket but in template handling, defining a route to next-generation B-family polymerase inhibitors.

**Graphical Abstract:** 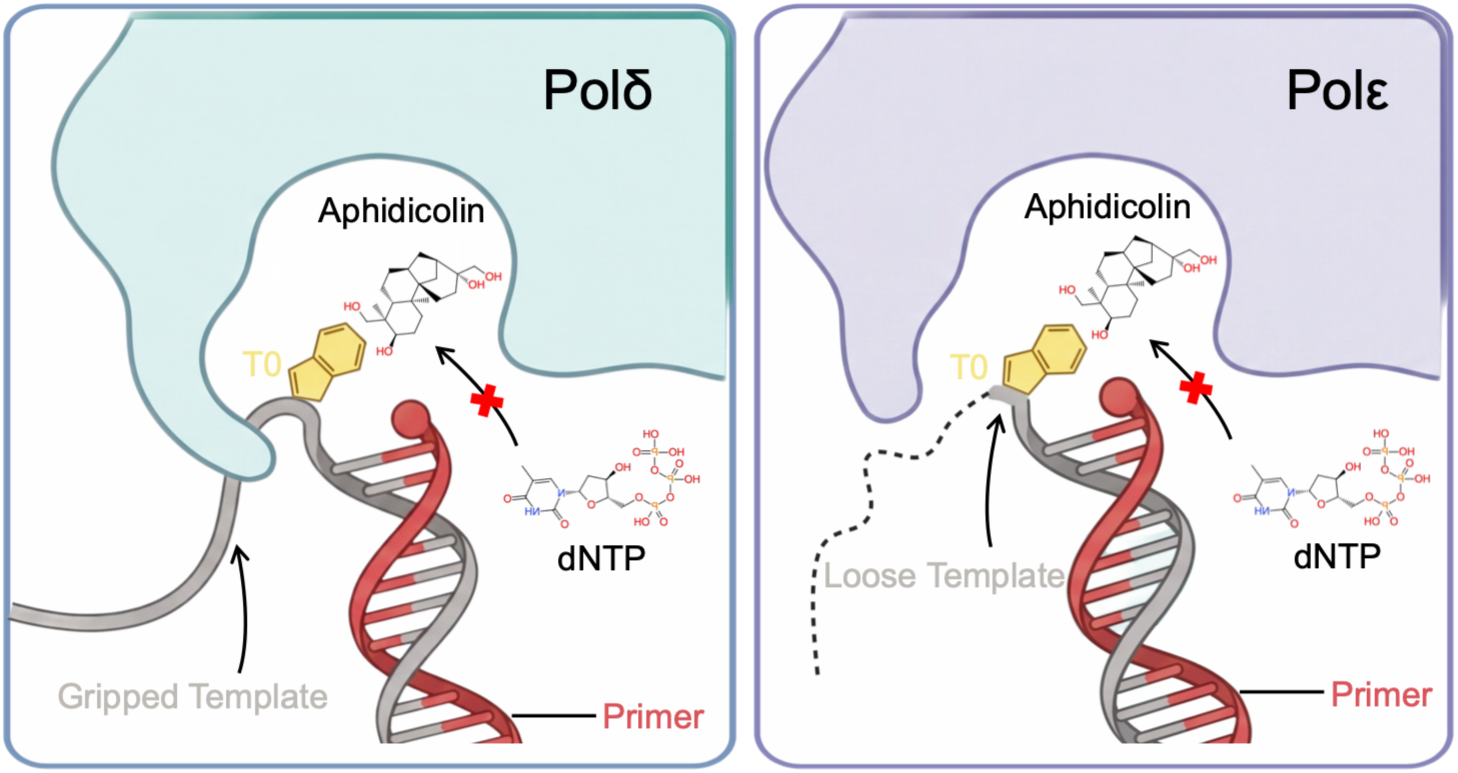

## Introduction

DNA replication in eukaryotes depends on highly accurate and processive replicative polymerases to duplicate billions of base pairs once per cell cycle while preserving genome integrity. At replication forks, the three B-family polymerases divide this task: Pol α−primase synthesizes RNA–DNA primers that are extended processively by Pol δ and Pol ε, which replicate the lagging and leading strands, respectively, in coordination with helicase-driven unwinding (1–3) . A central feature enabling rapid fork movement is polymerase processivity—the ability to incorporate long tracts of nucleotides without dissociating from the primer-template (PT) DNA (4–6) . For Pol δ and Pol ε, high processivity is conferred by the PCNA clamp, which tethers the polymerases to PT DNA and maintains stable engagement during rapid genome duplication (7–9) .

Aphidicolin has long served as a powerful probe of this process. First described as a natural product from the fungus *Cephalosporium aphidicola*, aphidicolin was initially recognized for its potent inhibition of cellular DNA synthesis and antiviral activity, most notably against herpes simplex virus (10, 11). These early observations quickly established aphidicolin as both a bioactive natural compound and a mechanistic tool. By transiently slowing replicative polymerase activity, aphidicolin became widely used to synchronize cells at the G1/S boundary and to induce controlled replication stress without broadly poisoning DNA metabolism (12, 13).

Classic kinetic studies showed that aphidicolin inhibits eukaryotic replicative DNA polymerases by interfering at the incoming dNTP site, most notably through strong competition against dCTP incorporation in Pol α-driven synthesis (14). Importantly, aphidicolin preferentially targets replicative polymerase activity, strongly inhibiting Pol α-dependent DNA synthesis while leaving Pol β and Pol γ largely insensitive, supporting its widespread use as a replication-focused inhibitor in cell biology (15). Extending beyond Pol α, mechanistic studies on Pol ε demonstrated that aphidicolin can compete with each of the four dNTPs for binding to a Pol ε-DNA complex and can bind with similar affinity independent of the next templating base, highlighting a binding mode that is not simply a nucleotide mimic but rather exploits conserved architectural features of the B-family active site (16).

Structural insight into this mode of inhibition came from a crystal structure of the catalytic core of human Pol α in complex with a primed template and aphidicolin, which placed the inhibitor near the incoming nucleotide site and showed that its binding induces flipping of the templating guanine, preventing the geometry required for catalysis (17). While informative, this structure captures only the minimal catalytic core of Pol α and therefore leaves unresolved how aphidicolin engages the multi-subunit, PCNA-tethered holoenzymes that actually carry out processive DNA synthesis.

Indeed, important gaps remain. It is unknown whether the mechanism defined on the minimal Pol α catalytic core extends to the multi-subunit, PCNA-tethered holoenzymes that carry out processive DNA synthesis, whose accessory subunits shape DNA engagement and clamp contacts (7–9). It is equally unexplained why the replicative polymerases differ in aphidicolin sensitivity: Pol α and Pol δ are inhibited 6- to 10-fold more potently than Pol ε (18), even though the inhibitor targets a conserved active-site pocket in all three enzymes.

Here, we employed Cryo-EM to resolve structures of human Pol δ and Pol ε holoenzymes bound to aphidicolin and defined how this canonical replication inhibitor arrests processive DNA synthesis in the PCNA-tethered, multi-subunit context most relevant to the replisome. By comparing inhibitor engagement, catalytic-site configurations, and DNA templating geometry across the two replicative holoenzymes, these structures provide a unified and polymerase-specific view of aphidicolin action, linking decades of kinetics and cell biology to an architectural mechanism for how processive synthesis is halted at eukaryotic replication forks.

## Materials and Methods

### Protein expression and purification

Wild-type human DNA polymerase δ (Pol δ WT) was produced using a previously established protocol (7). In short, a MultiBac™ plasmid carrying the genes for all four Pol δ subunits was transformed into DH10MultiBac™ cells to produce bacmid DNA. The resulting bacmid was then transfected into Sf9 insect cells with FuGENE® HD transfection reagent according to the manufacturer’s instructions, generating recombinant baculovirus. The virus was amplified through two successive passages to yield a high-titer P3 stock. Protein expression was carried out by infecting 4 liters of Sf9 cells in suspension culture (at a density of 2 × 10⁶ cells/mL) with the P3 viral stock, followed by incubation for 70 hours. After harvesting, cells were lysed and the clarified extract was initially purified by nickel-affinity chromatography on a HisTrap HP column (Cytiva) under low-salt buffer conditions. The eluate was further resolved by anion-exchange chromatography using a Mono Q column (Cytiva). Fractions confirmed to contain the complete four-subunit Pol δ complex were pooled and subjected to final polishing by size-exclusion chromatography on a Superdex 200 column (Cytiva). The purified Pol δ was concentrated, combined from peak fractions, aliquoted, snap-frozen in liquid nitrogen, and stored at −80 °C.

Human Pol ε was produced and purified following an established protocol (19). In brief, 4 liters of Hi5 cells in suspension culture (2 × 10⁶ cells/mL) were infected with high-titer P3 baculovirus carrying the Pol ε construct. After 72 hours of incubation, cells were pelleted, lysed, and the clarified supernatant was applied to a 5 mL HisTrap HP column (Cytiva). Bound material was eluted using an imidazole gradient. The Pol ε-containing eluate was immediately loaded onto a 5 mL Strep-Tactin XT column (Cytiva) and eluted with biotin. Relevant fractions were pooled and digested overnight with TEV protease to cleave affinity tags. The sample was then passed over a 1 mL Mono Q 5/50 GL anion-exchange column (Cytiva) and eluted with a gradient to 1 M NaCl. Peak fractions were concentrated to ∼1 mL and further purified by size-exclusion chromatography on a HiLoad 16/600 Superdex 200 pg column (Cytiva). Pure Pol ε fractions were combined, aliquoted, snap-frozen in liquid nitrogen, and stored at −80 °C.

The full-length human PCNA gene (UniProt: P12004) was inserted into the MCS1 site of pETDuet-1 (Novagen) with an ampicillin-resistance marker to produce an N-terminally 6×His-tagged protein. PCNA was expressed and purified as described previously (7). In brief, expression was performed in 2YT medium containing ampicillin. Cells were cultured at 37 °C until the OD_600_ reached ∼1.2, then induced with 0.5 mM IPTG and grown overnight at 16 °C for 19 hours. After harvesting by centrifugation, the cell pellet was resuspended in lysis buffer and disrupted by lysozyme digestion followed by sonication. All purification steps were conducted at 4 °C. Debris was removed by high-speed centrifugation, and the soluble fraction was applied to a pre-equilibrated HisTrap HP column (Cytiva). After thorough washing, His-tagged PCNA was eluted with an increasing imidazole gradient. The eluted material was further separated by anion-exchange chromatography on a HiTrap Q HP column (Cytiva), followed by size-exclusion chromatography on a Superdex 200 column (Cytiva) in storage buffer. Pure PCNA fractions were combined, concentrated, aliquoted, snap-frozen in liquid nitrogen, and stored at −80 °C.

### PT DNA substrate

An equimolar amount of a 25-nt primer (5’ AGCTATGACCATGATTACGAATTGC_dd_ 3’) was annealed to a 38-nt template (5’ CTGCACGAATTAAGCAATTCGTAATCATGGTCATAGCT 3’) by heating at 95 °C for 2 min in 10 mM HEPES, pH 7.5, and 50 mM NaCl, then slowly cooled to room temperature. The annealed substrate was stored at -20 °C until use.

### Cryo-EM sample preparation

Pol δ, PCNA, and PT DNA were combined at equimolar ratios to a final concentration of 0.5 µM each in buffer containing 20 mM HEPES pH 7.5, 150 mM NaCl, 5 mM CaCl₂, and 1 mM DTT. The mixture was incubated on ice for 10 min. Aphidicolin was then added to 250 µM, and incubation continued for an additional 20 min to assemble the complex. 3 µL aliquot of the reaction was applied to glow-discharged (40 mAmp, 5 mins) UltrAuFoil R1.2/1.3 300-mesh grids that had been precoated with a thin layer of graphene oxide. Grids were blotted and plunge-frozen in liquid ethane using a Thermo Fisher FEI Vitrobot set to 4 °C and 100% humidity. Frozen grids were stored in liquid nitrogen until cryo-EM data collection.

In a similar manner, Pol ε, PCNA, and PT DNA were mixed in a molar ratio of 1:2.3:1.1 resulting in final concentrations of 3 µM Pol ε, 6.8 µM PCNA, and 3.4 µM DNA in buffer containing 20 mM HEPES pH 7.5, 200 mM NaCl, 2 mM CaCl₂, and 1 mM DTT. Aphidicolin was added to a final concentration of 250 µM, and the reaction was incubated for 10 min at room temperature, followed by 2 h on ice to promote complex formation. 3 µL sample was applied to Quantifoil Cu R2/1 300-mesh grids (coated with a 2 nm carbon layer) that had been glow-discharged at 10 mA for 10 s. Grids were blotted and plunge-frozen in liquid ethane using a Thermo Fisher FEI Vitrobot operated at 4 °C and 100% humidity. Vitrified grids were stored in liquid nitrogen until cryo-EM data collection.

### Cryo-EM data collection

All cryo-EM data were collected on a 300 kV Titan Krios G4 (Thermo Fisher Scientific) equipped with a Falcon 4 direct electron detector at the Imaging and Characterization Core Lab, King Abdullah University of Science and Technology (KAUST). Data were acquired in electron counting mode using EPU (Thermo Fisher Scientific) and recorded as dose-fractionated movies in EER format. For both Pol δ (Dataset 1) and Pol ε (Dataset 2) holoenzyme–inhibitor complexes, movies were collected at a nominal magnification of 165,000×, corresponding to a calibrated pixel size of 0.76 Å pixel⁻¹. Dataset 1 was acquired with a total electron dose of 42.62 e⁻ Å⁻² and a defocus range of −3.0 to −1.0 µm, while the Dataset 2 was collected with a total dose of 43.34 e⁻ Å⁻² and a defocus range of −2.0 to −1.0 µm.

### Cryo-EM data processing

For both datasets (12,002 movies for Dataset 1 and 34,984 movies for Dataset 2, recorded at 0.76 Å/pixel), the acquired movies were subjected to beam-induced motion correction using the built-in motion correction implementation in RELION (20, 21) . Dose weighting was applied during motion correction. Contrast transfer function (CTF) parameters were subsequently estimated using Patch CTF estimation as implemented in cryoSPARC (22) . For Dataset 1, particle picking was initially performed using the blob-based auto-picking approach in cryoSPARC to generate a preliminary particle set of 6,418,570 particles, extracted in a 400-pixel box. These particles were extracted and subjected to three rounds of 2D classification to remove obvious contaminants and poorly aligned particles, retaining 90,048 particles. The resulting particle subsets were then used to train a Topaz neural network model (23) , which was applied for Topaz-based particle picking to enrich for true particle projections; for Dataset 2, Topaz picking was applied directly to the full micrograph set, yielding 8,264,972 particles in a 400-pixel box. Particles obtained from Topaz picking were re-extracted and further cleaned through iterative rounds of 2D classification, leaving 262,654 particles after two rounds (Dataset 1) and 345,955 particles after three rounds (Dataset 2). Selected particle stacks were subsequently subjected to ab initio reconstruction in cryoSPARC to generate initial models; for Dataset 2, two further rounds of 3D classification isolated a homogeneous subset of 123,747 particles. These models were refined using non-uniform refinement in cryoSPARC, giving a 2.9 Å and 3.1 Å reconstructions for Dataset 1 and Dataset 2. For final refinement and correction of dataset anisotropy, the refined particle stacks were exported to RELION and subjected to 3D auto-refinement. SpIsoNet (24) was applied to correct for preferred orientation–induced anisotropy and misalignment effects, resulting in improved isotropy of the reconstructed maps, at 3.1 Å (Dataset 1) and 3.2 Å (Dataset 2). A final spIsoNet anisotropy correction during post-processing yielded the Dataset 1 consensus map at 3.0 Å, while a final round of non-uniform refinement in cryoSPARC yielded the Dataset 2 map at 3.1 Å. For Dataset 1, a soft mask encompassing PCNA was used for local refinement, which reached 3.5 Å; this map was combined with the consensus reconstruction to produce the composite map used for model building. All final maps were post-processed with EMReady2 (25) prior to model building. All resolutions are reported at the FSC = 0.143 criterion.

### Model building and visualizations

The model of the Pol δ (PDB 6TNY) or Pol ε holoenzymes (PDB 9B8S) were docked into the refined cryo-EM maps along with aphidicolin. The models were refined using iterative ISOLDE flexible fitting and Phenix real space refine (26–29) . Phenix Cryo-EM Standard validation was performed to finalize the structure. Finally, protein-ligand interactions were computed using PLIP 2025 (30).

### Primer extension assay

Primer extension assays were performed using Cy5-labelled primer-annealed M13 ssDNA as the substrate. Reactions were assembled in a final volume of 10 µl in primer-extension buffer containing 50 mM HEPES (pH 7.5), 80 mM KCl, 12 mM magnesium acetate, 2 mM MgCl₂, 5 mM DTT and 0.1 mg/mL BSA. Each complete reaction contained 150 nM PCNA, 40 nM ΔN-RFC (for Pol δ) or CTF18-RFC (for Pol ε), 300 nM RPA, and 60 nM Pol δ or Pol ε, together with 5 nM of Cy5-primed M13 substrate and 1 mM ATP. 500 µM of each of dATP, dGTP and dCTP were added during reaction assembly. To assess inhibition of DNA synthesis, Pol δ and Pol ε were tested in the presence of increasing concentrations of inhibitor (1, 5, 10, 25, 50, 75 and 100 µM), with uninhibited reactions included as controls. The inhibitor was prepared as a 5 mM stock in DMSO and diluted to the required working concentrations in primer-extension buffer. DNA synthesis was initiated by the simultaneous addition of 500 µM dTTP, and the indicated concentration of inhibitor and reactions were incubated at 30°C for 15 min. Reactions were terminated by adding 4 µl of 100 mM EDTA followed by 2 µl of 5X loading buffer. The reaction products were resolved on a 1% alkaline agarose gel and Cy5-labelled DNA products were visualized using a Typhoon fluorescence scanner.

To quantify polymerase inhibition, a median length analysis was performed on the extended DNA products, as previously described (31–33). The median synthesis length for a given lane is defined as the absolute nucleotide length at which 50% of the total integrated signal intensity is reached within a defined lane boundary. Background signal was subtracted prior to integration using the Sub-alone and Pol-alone control lanes. Migration distances along the gel were converted to absolute nucleotide lengths by interpolating against a standard curve generated from the known marker sizes (3, 4, 5, 6, and 8 Kb). To account for assay-specific background, the integration areas used to calculate the median lengths were distinct for each polymerase. Pol δ lanes were integrated from ∼350 nt to 9000 nt. For Pol ε, integration was restricted to the region between ∼850 nt and 9000 nt to exclude the bottom persistent band and its shadow (running from ∼200 nt to ∼700 nt). The calculated median lengths (*L*) were plotted as a function of aphidicolin concentration ([*I*]). Because aphidicolin acts as a competitive inhibitor of dNTPs, the data were fit to a hyperbolic decay model that reflects the standard dose-response kinetics foundational to the Cheng–Prusoff relationship (34). The fit utilized the following equation:

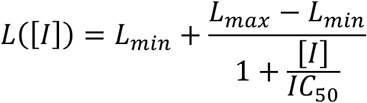

where *Lmax* is the maximum median length in the absence of inhibitor, *Lmin* is the minimum baseline length at saturating inhibitor concentrations, and *IC50* is the apparent half-maximal inhibitory concentration, representing the concentration of aphidicolin required to reduce the median product length to exactly halfway between *Lmax* and *Lmin*.

## Results

### Overall architectures of aphidicolin-bound human Pol δ and Pol ε holoenzymes

To visualize how aphidicolin inhibits the replicative polymerase holoenzymes, we reconstituted human Pol δ and Pol ε with PCNA, a 25/38-nt PT DNA carrying a 3’-dideoxy-terminated primer, in the presence of saturating aphidicolin and CaCl_2_. We then determined their structures by single-particle cryo-EM to 3.0 Å and 3.1 Å resolution, respectively (Figure 1, Figure S1-2).

**Figure 1:**
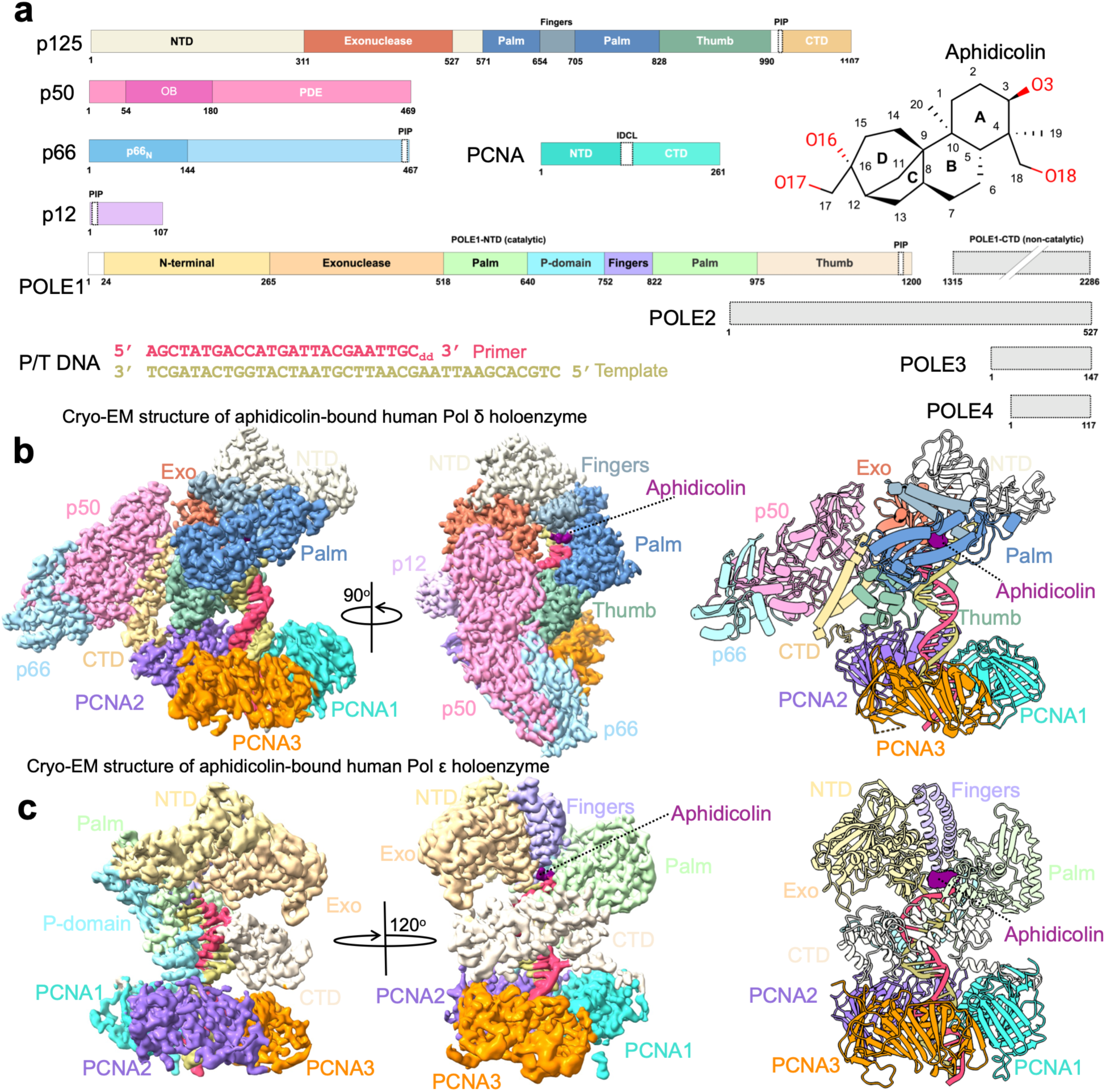
Cryo-EM structures of aphidicolin-bound human Pol δ and Pol ε holoenzymes. **(a)** Domain architecture of the Pol δ subunits p125, p50, p66, and p12, of PCNA, and of the Pol ε subunits POLE1–POLE4, with the catalytic POLE1 NTD (residues 1–1230) and non-catalytic CTD (1315–2286) indicated. PIP boxes and the interdomain connecting loop (IDCL) are marked. Right, chemical structure of aphidicolin with the four fused rings (A–D) and hydroxyl groups O3, O16, O17, and O18 highlighted. Below, the primer-template (P/T) DNA substrate used for cryo-EM; the primer 3′-terminal dideoxycytosine (ddC) blocks further extension. (**b)** Cryo-EM density (left, middle) and ribbon model (right) of the aphidicolin-bound Pol δ holoenzyme (p125, p50, p66, p12, PCNA). Aphidicolin (purple) is bound in the polymerase active site. (**c)** As in b, for the aphidicolin-bound Pol ε holoenzyme (POLE1 NTD, PCNA), with a 120° rotation between the density views.

The Pol δ reconstruction resolves all four subunits of the polymerase, the catalytic p125 and the regulatory p50, p66, and p12 subunits, engaging a P/T DNA duplex that threads through the PCNA trimer (Figure 1a-b). The catalytic p125 subunit adopts the canonical B-family right-hand fold, with its palm, fingers, and thumb subdomains arranged around the primer terminus, while p66 contacts p50 and p12 bridges p125 and p50 to complete the tetrameric assembly. The polymerase sits on one face of the PCNA ring through the p125 thumb and C-terminal domain. A well-defined density for aphidicolin is observed in the active-site cleft, wedged between the fingers and palm subdomains immediately adjacent to the primer 3’-end (Figure 1b).

The Pol ε reconstruction resolves the catalytic NTD of POLE1 (residues 24-1200), encompassing the exonuclease, palm, fingers, and thumb subdomains, engaged with the PT DNA that threads through the PCNA trimer (Figure 1c). The accessory subunits POLE2, POLE3, and POLE4 are not resolved, consistent with their flexible tethering observed in prior cryo-EM studies of Pol ε (8, 9). The catalytic NTD sits on one face of PCNA through its P-domain, thumb and C-terminal region. As in Pol δ, aphidicolin is unambiguously resolved at the fingers-palm interface, positioned immediately above the primer 3’-end (Figure 1c). Overall, both holoenzymes adopt the canonical organization of their respective nucleotide-bound ternary complexes (7–9), with aphidicolin positioned in the incoming-nucleotide binding site.

### Aphidicolin blocks nucleotide incorporation by remodeling pre-insertion sites

The high-resolution density in both reconstructions allowed unambiguous placement of aphidicolin and the surrounding DNA in the active-site cleft (Figure 1, Figure 2a, Figure 3a). In both holoenzymes, the inhibitor is placed between the fingers and palm subdomains directly above the primer 3’-end and occupies the volume that the incoming dNTP would otherwise fill. The PT DNA is tracked continuously into the active-site cleft, with the primer terminus positioned at the active site ready for nucleotide incorporation. At the incoming dNTP insertion site, the templating base (T0) is flipped out of the helical stack by aphidicolin, disrupting the pre-insertion geometry required for catalysis. (Figure 2a, Figure 3a). The templating base at T0 is an adenine in both structures. To test whether the flipped-out pose depends on base identity, we substituted each of the four bases in turn at T0 in both structures; none produced atomic clashes with the surrounding protein or aphidicolin (closest heavy-atom contacts ≥ 2.85 Å; Figure S3). The extruded-base pose therefore tolerates any templating base, providing a structural rationale for the base-independent inhibition of Pol ε by aphidicolin (16) .

**Figure 2:**
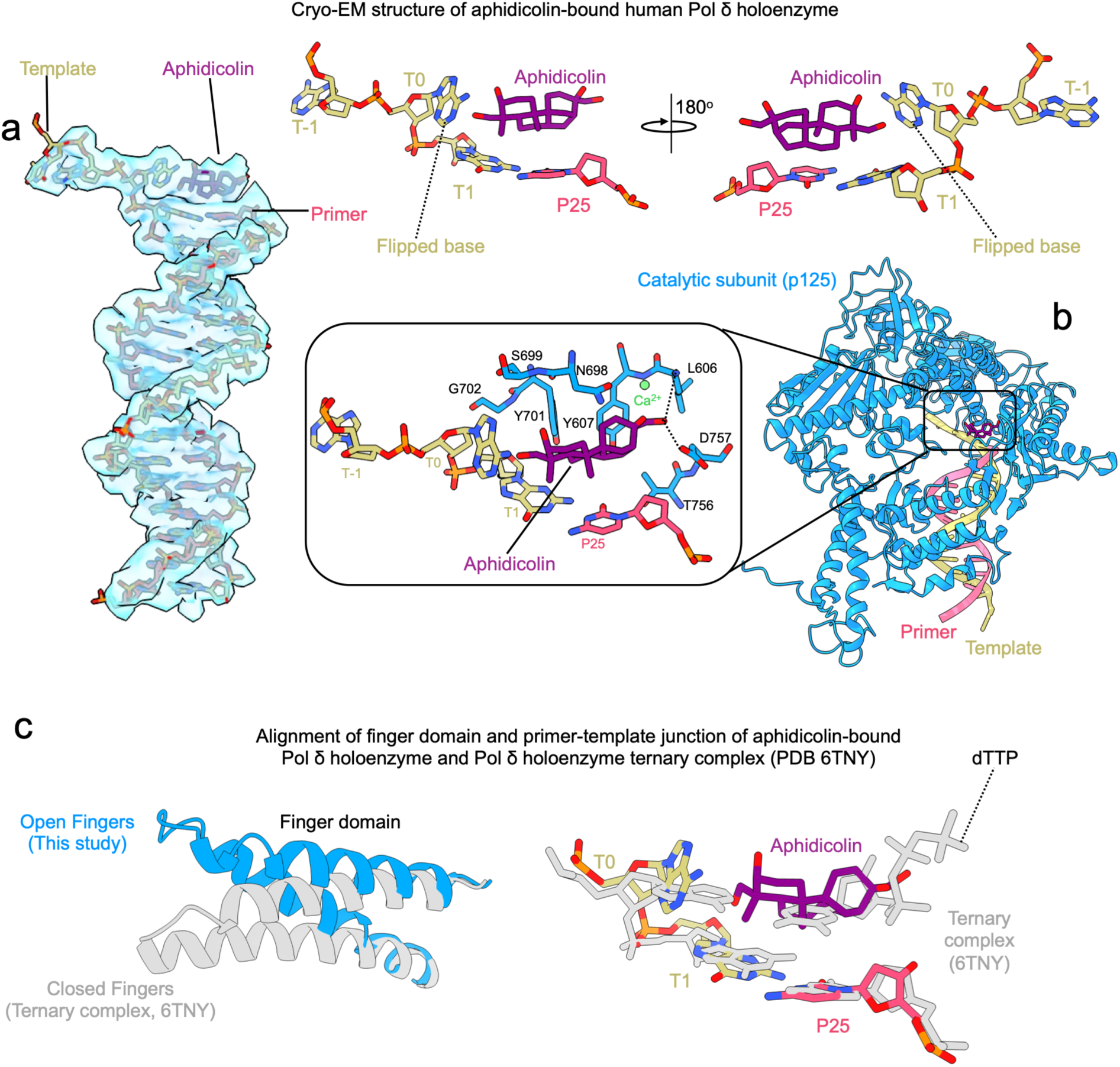
Analysis of the aphidicolin-bound Pol δ active site. **(a)** Cryo-EM density of the primer-template junction in the aphidicolin-bound Pol δ holoenzyme. Close-ups (two orientations related by 180°) show aphidicolin bound at the templating (T0) position opposite the primer 3′ terminus (P25), with the template base at T0 flipped out of the stacking register. **(b)** Active-site close-up of the catalytic subunit p125. Aphidicolin is wedged between the palm (L606, Y607, T756, D757) and the primer terminus (P25). Hydrogen bonds involving D757 and L606 backbone are shown as dashed lines. Fingers residues N698, S699, Y701 and G702 line the pocket but make no contact with the inhibitor. The inhibitor displaces the templating base and sterically replaces the incoming nucleotide. **(c)** Structural alignment of the finger domain (left) and primer-template junction (right) of the aphidicolin-bound Pol δ holoenzyme (this study) with the dTTP-bound ternary complex (PDB 6TNY) (7). The fingers remain in an open conformation upon aphidicolin binding, in contrast to the closed conformation of the catalytically active ternary complex.

**Figure 3:**
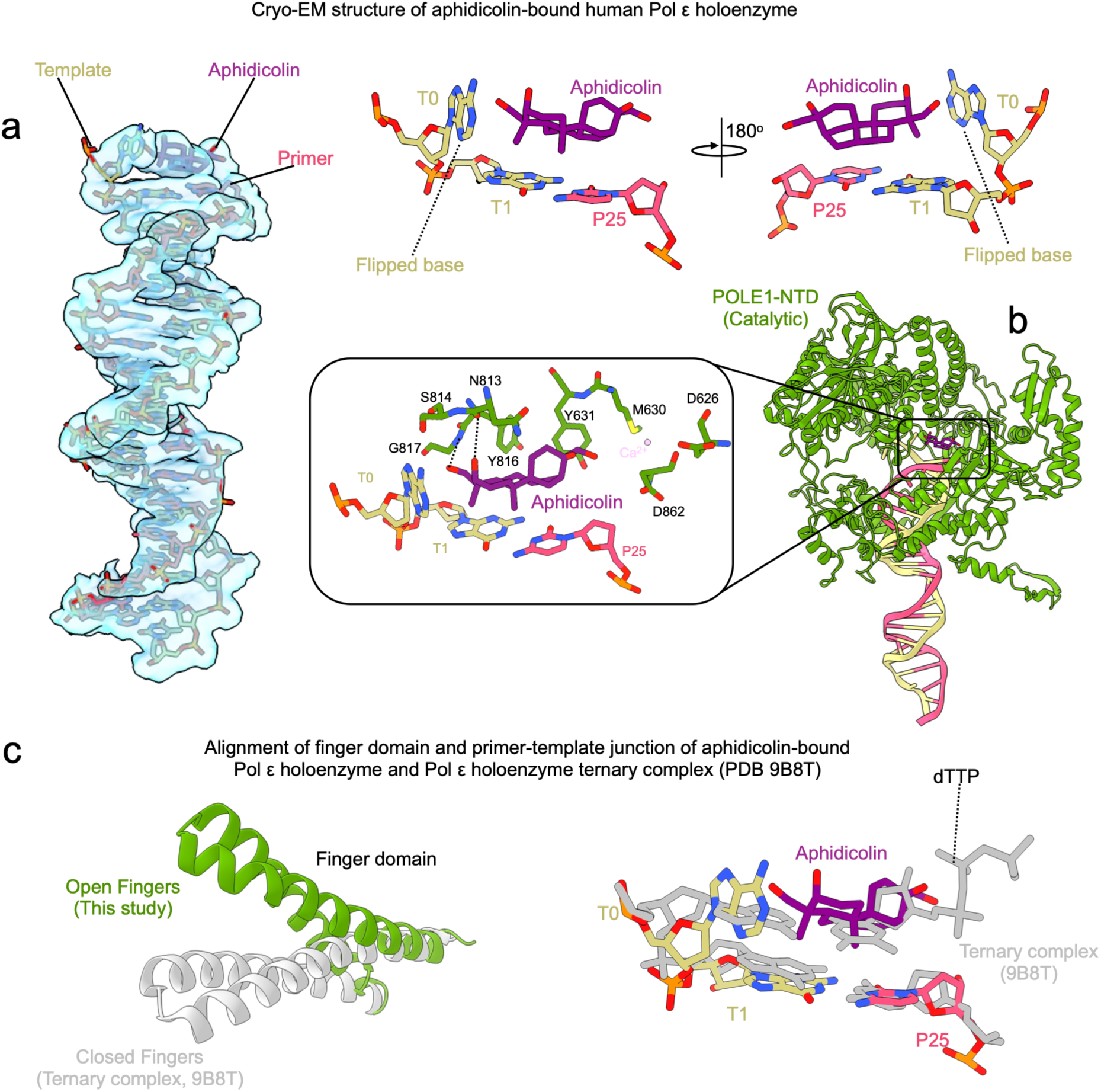
Analysis of the aphidicolin-bound Pol ε active site. **(a)** Cryo-EM density of the primer-template junction in the aphidicolin-bound Pol ε holoenzyme. Close-ups (two orientations related by 180°) show aphidicolin bound at the T0 position above P25, with the T0 template base flipped out, mirroring the Pol δ configuration. **(b)** Active-site close-up of the catalytic POLE1 NTD. Aphidicolin is wedged between the finger domain (N813, Y816, G817) and the palm domain (Y631). Hydrogen bonds involving N813 and G817 backbone are shown as dashed lines. S814, D626, M630 and D862 line the pocket but make no contact with the inhibitor. (**c)** Structural alignment of the finger domain (left) and primer-template junction (right) of the aphidicolin-bound Pol ε holoenzyme (this study) with the dTTP-bound ternary complex (PDB 9B8T) (9) . As in Pol δ, the Pol ε fingers remain open upon aphidicolin binding.

In the Pol δ active site, aphidicolin is coordinated by p125 palm domain residues L606, Y607, and D757 (Figure 2b). The tetracyclic core adopts a chair-like conformation and stacks directly above the P25–T1 primer-terminal base pair. One face of the scaffold is stabilized by hydrophobic contacts with Y607 while a D-ring hydroxyl form polar interactions with D757 and the backbone of L606. These interactions position aphidicolin directly above the primer terminus, blocking the incoming nucleotide site and preventing the flipped T0 base from returning to a productive templating position. A similar arrangement is observed in Pol ε. Aphidicolin is coordinated by Y631 of the palm domain together with N813, Y816 and G817 of the fingers domain (Figure 3b). The aphidicolin tetracyclic core adopts the same conformation above the P25-T1 base pair, stacking with the primer-template terminus rather than pairing the flipped T0 base. Hydrogen bonds involving the backbone of N813 and G817, together with hydrophobic contacts with Y631 and Y816, stabilize aphidicolin binding. Although the specific hydrogen bonds differ between the two polymerases, equivalent residues occupy every contact position in both enzymes and could form the reported interactions. This shared mode of interaction, in which aphidicolin displaces the templating base and blocks the incoming nucleotide site, indicates that it exploits conserved features of the B-family polymerase pre-insertion site rather than enzyme-specific chemistry.

Comparison of each aphidicolin-bound holoenzyme with the corresponding nucleotide-bound ternary complex (7, 9) revealed two key features (Figure 2c, Figure 3c). First, the fingers domain adopts an open conformation in the inhibited state relative to the closed fingers seen in the ternary complexes, showing that aphidicolin traps the polymerase in a pre-catalytic open-fingers state incompatible with nucleotide alignment and catalytic closure. Second, the superposition of the ternary-complex dNTPs onto the inhibited maps confirms that aphidicolin occupies the same volume as the incoming nucleotide but displaces the templating base rather than pairing with it, distinguishing it from a classical nucleotide mimic. Together, these observations define a shared inhibitory mechanism in which aphidicolin occupies the incoming-nucleotide pocket, pushes the templating base out of the duplex, and locks the fingers domain open, preventing the conformational closure required for phosphodiester-bond formation.

### Differences in template gripping between Pol δ and Pol ε

The two holoenzymes handle the template strand beyond the inhibitor-bound templating base in markedly different ways. In the aphidicolin-bound Pol δ structure, two template bases beyond T0 (T-1 and T-2) are resolved and anchored by bona fide protein contacts: the side chain of Q552 and the backbone of G550 and Q551 form hydrogen bonds with the phosphate backbone of T-1 and T-2, and the side chain of R549 makes an additional salt bridge with the backbone phosphates (Figure 4a,b). The same contacts are present in the dTTP-bound Pol δ holoenzyme (7), indicating that the grip is maintained independently of the nucleotide-bound state. Pol ε shows the opposite behavior. No density beyond T0 is observed in the aphidicolin-bound structure. Density beyond T0 is resolved only in the active, dTTP-bound state with closed fingers (PDB 9B8T), where the template strand is anchored by the fingers-domain residue R821 together with the EXO-domain residues R494, G496, S497, G498, and T499 (9). In the inactive, open-fingers state (PDB 9B8S), this density is absent, and the anchoring residues retract from the template beyond T0 and reposition at the primer-template junction (Figure S4c,d). Pol ε therefore anchors the template beyond T0 only when its fingers close.

**Figure 4:**
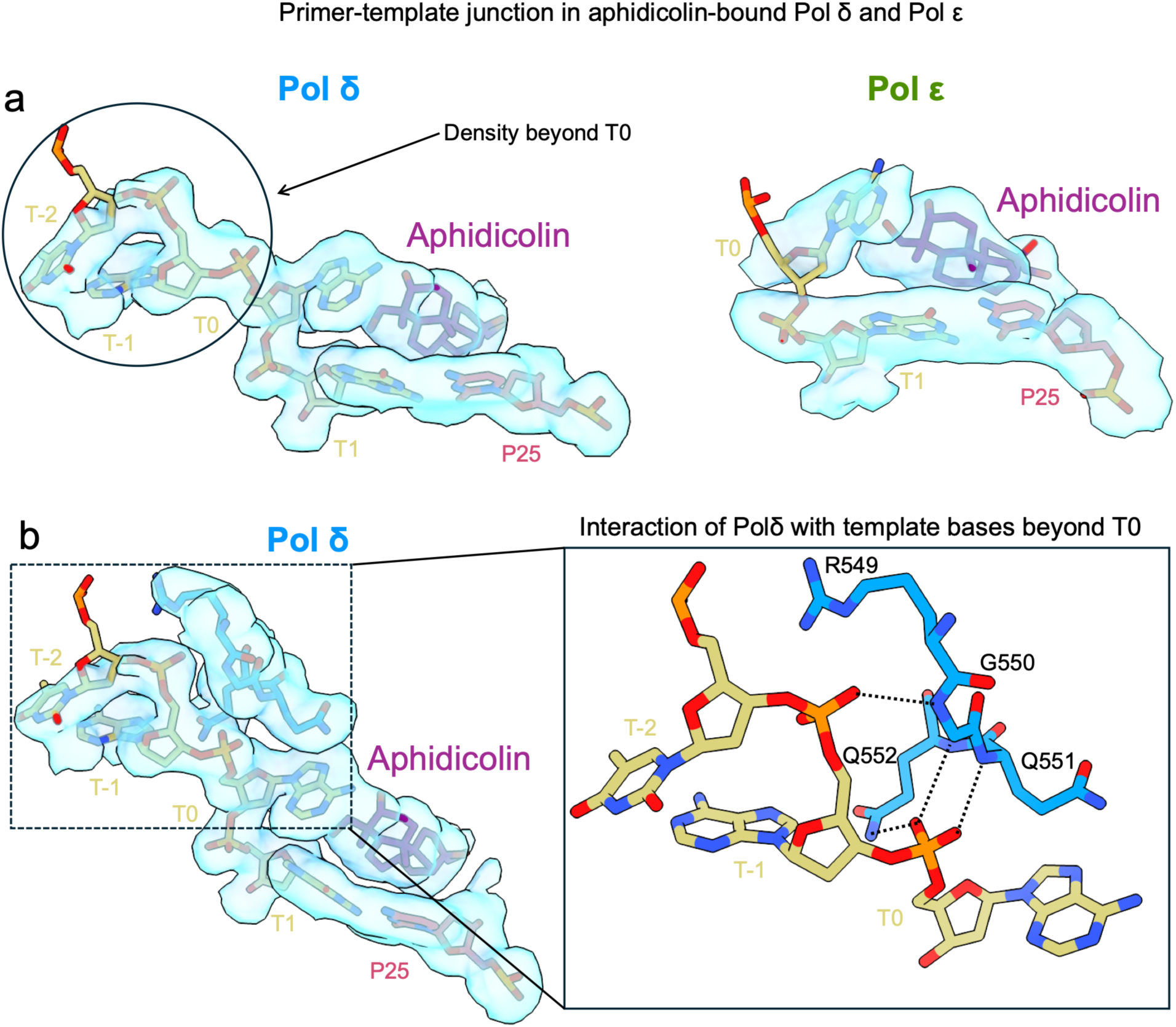
Comparison of template-strand density beyond the templating base in aphidicolin-bound Pol δ and Pol ε. **(a)** Cryo-EM density at the primer-template junction shows ordered density beyond the templating base (T-1, T-2) in Pol δ, whereas in Pol ε the template beyond T0 is disordered. **(b)** In Pol δ, R549, G550, Q551, and Q552 contact the phosphate backbone of template bases T-2 and T-1, anchoring the template strand upstream of the aphidicolin-bound templating site. No equivalent contacts are present in Pol ε.

### The grip accounts for differential aphidicolin sensitivity

A difference in template handling suggests a mechanism for a long-standing but unexplained observation: aphidicolin inhibits Pol δ with 6-to-10 fold higher potency than Pol ε in calf thymus enzymes (18). We hypothesized that the grip accounts for this selectivity — a rigidly gripped template remains locked in the inhibitory register even when the fingers are open, whereas the loosely anchored Pol ε template can disengage and re-bind productively, escaping the aphidicolin trap. This predicts that the human PCNA-tethered holoenzymes, not previously compared, differ in aphidicolin sensitivity. To test this, we compared the two reconstituted holoenzymes in primer-extension assays on a primed M13mp18 template in the presence of their respective PCNA loading factors, RFC and CTF18-RFC, across increasing aphidicolin concentrations (0–100 µM) (Figure 5a). In the absence of inhibitor, both Pol δ and Pol ε generated long extension products of up to ∼8 kb, consistent with highly processive DNA synthesis by the PCNA-tethered holoenzymes. Aphidicolin titration revealed markedly different responses. For Pol δ, full-length products were sustained only up to 1 µM aphidicolin and were largely abolished at 25 µM, with only short primer-proximal fragments remaining at higher concentrations. In contrast, Pol ε retained full-length synthesis up to 10 µM and continued to produce long extension products across most of the concentration range tested. Quantification of median product lengths yielded hyperbolic inhibition curves with IC50 values of 3.91 ± 0.48 µM for Pol δ and 18.13 ± 8.99 µM for Pol ε (Figure 5b,c): Pol δ is approximately 4-5-fold more sensitive to aphidicolin than Pol ε, consistent with the calf thymus selectivity (18) and matching the prediction of the grip mechanism.

**Figure 5:**
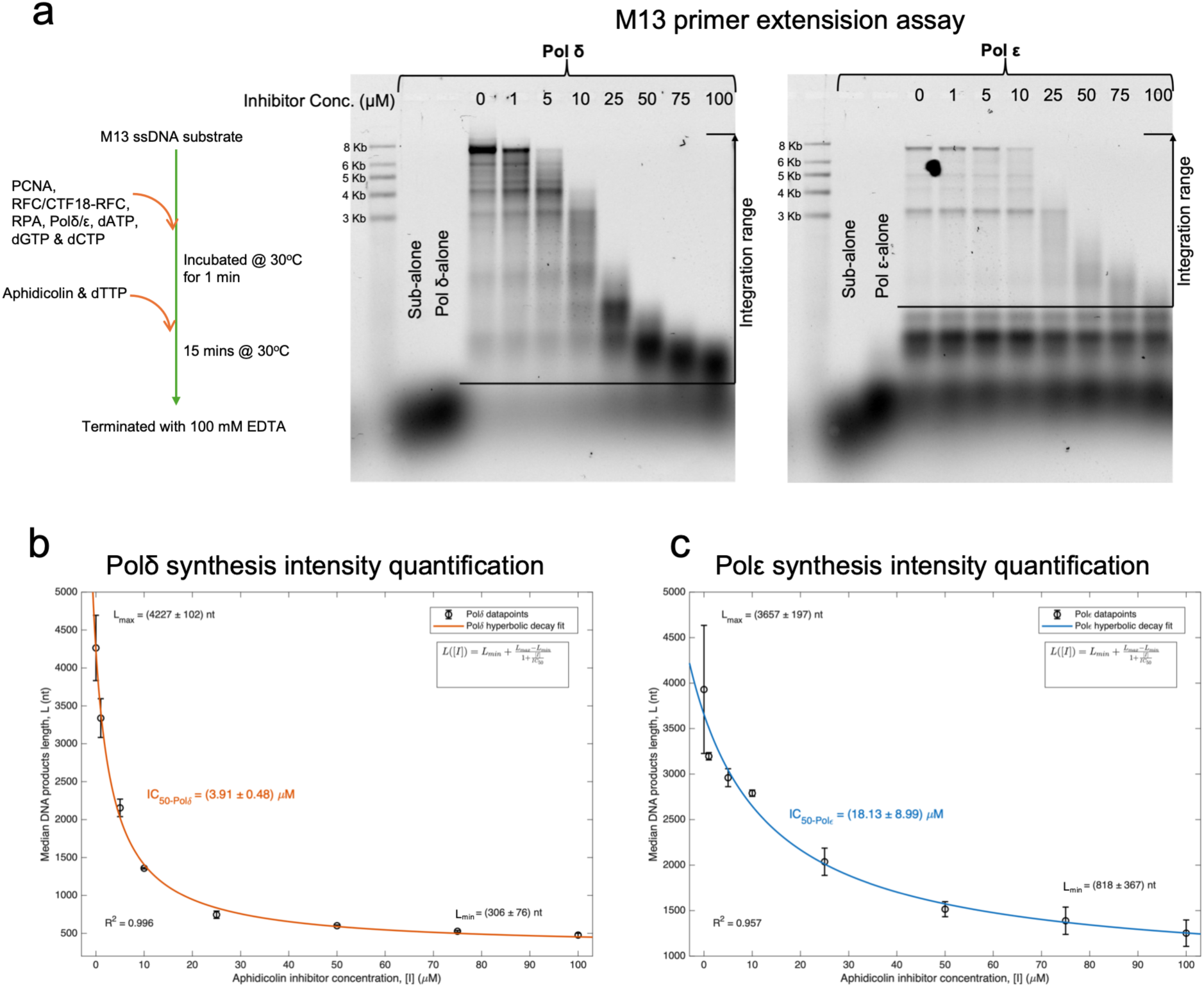
M13 primer extension of PCNA-tethered Pol δ and Pol ε holoenzymes across an aphidicolin titration. **(a)** Schematic of the primer-extension assay. Reactions containing the M13 ssDNA substrate, PCNA, RPA, Pol δ or Pol ε, the corresponding clamp loader (RFC or CTF18-RFC, respectively), and 500 µM each of dATP, dGTP and dCTP were pre-incubated at 30 °C for 1 min. Aphidicolin and dTTP were then added, and reactions were incubated at 30 °C for 15 min before termination with 100 mM EDTA. The assay was performed in triplicate. Right, gel showing primer-extension products generated at aphidicolin concentrations ranging from 0 to 100 µM. Size markers and the 25-nt primer are indicated. Brackets indicate the integration range used for median length quantification (described in the Materials and Methods section). **(b)** Aphidicolin dose-response curve for Pol δ. Absolute median DNA product length synthesized by Pol δ is plotted as a function of aphidicolin concentration. Median synthesis length is defined as the product length at which 50% of the total synthesis intensity within the defined integration range (∼350 to 9000 nt; indicated by brackets in the representative gels in panel a) is achieved at the fixed 15-minute timepoint. Data points are fit to a standard hyperbolic decay model linked to the Cheng–Prusoff relationship for competitive inhibition (*R*^2^ = 0.996), yielding an apparent *IC*_50_ of 3.91 ± 0.48 µM. Data points represent the mean of three independent experiments ± SEM, and the reported error for the *IC*_50_ represents one standard deviation. **(c)** Aphidicolin dose-response curve for Pol ε. Absolute median DNA product length synthesized by Pol ε is plotted as a function of aphidicolin concentration. Median synthesis length is determined as described in (b), except the lane integration range was restricted to approximately ∼850 to 9000 nt (indicated by brackets in panel a) to exclude the bottom persistent band and its associated shadow (running from ∼200 nt to ∼700 nt), which are present even at zero aphidicolin concentration. The data points are fit to the same hyperbolic decay model (*R*^2^ = 0.957), yielding an apparent *IC*_50_ of 18.13 ± 8.99 µM. Data points represent the mean of three independent experiments ± SEM, with the *IC*_50_ error representing one standard deviation.

## Discussion

### Aphidicolin arrests DNA synthesis by trapping a pre-insertion state

Aphidicolin is a potent, reversible inhibitor of eukaryotic DNA replication, yet its mechanism has remained difficult to reconcile with its chemistry. Early biochemical studies established that aphidicolin competes with dNTPs for binding to the polymerase–DNA complex, particularly with dCTP during Pol α-driven synthesis (35), and later work showed that inhibition proceeds through a stable polymerase–DNA–aphidicolin ternary complex rather than direct binding to the free enzyme (36). Subsequent studies on Pol ε showed that aphidicolin competes with all four dNTPs with little dependence on the identity of the templating base (16). This behavior is inconsistent with classical nucleotide mimicry, as aphidicolin is a tetracyclic diterpene that lacks structural similarity to dNTPs. The crystal structure of the Pol α catalytic core bound to a primed template and aphidicolin resolved this paradox: the inhibitor occupies the incoming dNTP pocket and rotates the templating base out of the helix (17) .

Whether this mechanism operates in the multi-subunit, PCNA-tethered holoenzymes that carry out processive synthesis — the context in which the drug acts at the replication fork — remained untested. Our cryo-EM structures show that it does. In both Pol δ and Pol ε holoenzymes, aphidicolin inserts between the fingers and palm, stacks above the primer-terminal base pair, and forces the templating base to rotate out of the helix, trapping the polymerase in an open-fingers, pre-catalytic state in which the template cannot present a Watson–Crick partner for nucleotide incorporation. Rather than mimicking an incoming nucleotide, aphidicolin blocks its binding site and disables templating by extruding the base itself. Modelling of all four bases at the flipped-out T0 position confirms that the pocket accommodates any templating base without clashes (Figure S3), providing a structural explanation for the ability of aphidicolin to compete with all four dNTPs and for its base-independent binding to Pol ε (16). Templating-base displacement coupled to an open-fingers trap is therefore a general mechanism of inhibition for B-family polymerases, conserved from the minimal Pol α core to the full PCNA-bound holoenzymes.

### Structural basis of aphidicolin sensitivity in human replicative polymerases

Although aphidicolin engages Pol δ and Pol ε in an almost identical manner, the two polymerases respond differently to the drug. In reconstituted PCNA-tethered reactions, Pol δ is inhibited at approximately 4-5-fold lower concentrations than Pol ε, with IC50 values of 3.91 µM and 18.13 µM, respectively (Figure 5). This difference has a biochemical precedent, as aphidicolin inhibits calf thymus Pol α and Pol δ with 6- to 10-fold higher potency than Pol ε (18). The inhibitor pocket itself, which is architecturally conserved in Pol α, Pol δ, and Pol ε, cannot explain this difference. Although the specific hydrogen bonds differ among the three polymerases, each contact position in one enzyme is occupied by an equivalent residue in the others that could form the reported interaction (Figure 6), and the observed pattern therefore likely represents a single snapshot of the inhibited state rather than the full range of contacts available. We found that the sensitivity emerges from how Pol δ and Pol ε polymerases handle the template strand beyond the templating base. In Pol δ, two template bases upstream of T0 remain ordered and are held by a dedicated grip formed by Q552, G550, Q551, and R549, contacts that persist in the nucleotide-bound state (7). In Pol ε, the template beyond T0 is only stabilized in the closed-finger state (9), through the finger residue R821 and a cluster of EXO-domain residues, and is released when the fingers open (9), leaving the upstream template flexibly anchored. We propose that this loose anchoring allows the template in Pol ε to disengage and re-bind productively, effectively escaping the aphidicolin trap, whereas the rigidly gripped template of Pol δ remains locked in the inhibitory register even with open fingers. The grip therefore acts as a sensitivity determinant that converts an otherwise identical inhibitor engagement into a polymerase-specific response.

**Figure 6:**
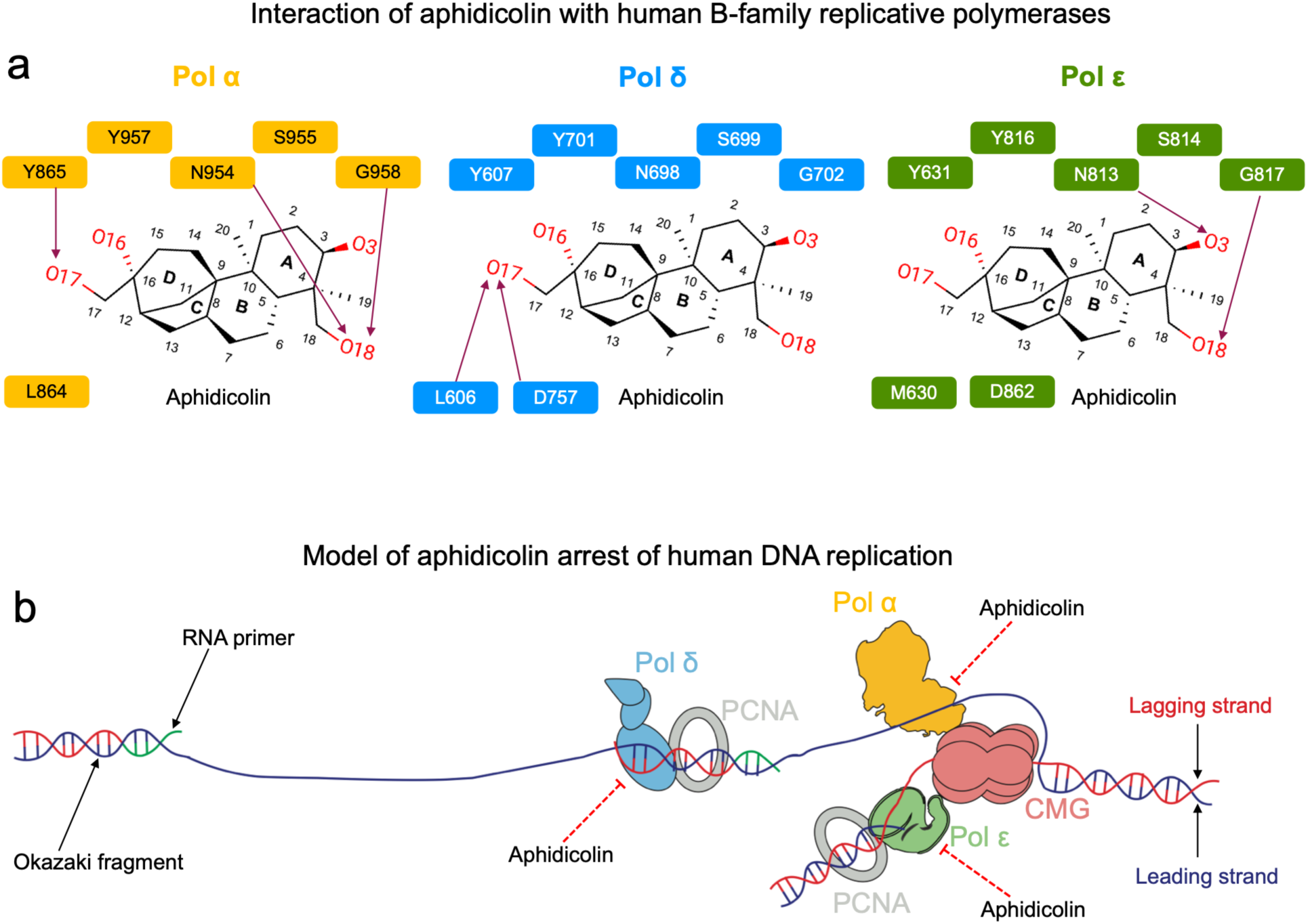
A conserved aphidicolin-binding mode and a model for aphidicolin arrest of human DNA replication. **a)** Interaction map of aphidicolin with the active sites of human B-family replicative polymerases, shown for Pol α (yellow) (17), Pol δ (blue), and Pol ε (green). Residues contacting the O3, O16, O17, and O18 hydroxyls and the ring system are indicated on the aphidicolin scaffold. (**b)** Model of aphidicolin arrest of human DNA replication at the replication fork. Aphidicolin binds the templating-base pocket of all three B-family polymerases on the leading strand (Pol ε), the lagging strand (Pol δ), and Okazaki-fragment priming (Pol α), stalling processive synthesis by PCNA-tethered holoenzymes and the CMG helicase-coupled replisome.

### Implications for replication stress

The differential sensitivity predicts an ordered response to aphidicolin at the replication fork rather than a uniform slowdown. Pol δ is the more sensitive of the two replicases (Figure 5), and Pol α, which synthesizes the primer of every Okazaki fragment, is at least as sensitive in vitro (18) . The lagging strand is therefore copied by two drug-sensitive polymerases acting in succession. Lagging-strand synthesis should slow first, and delayed Okazaki fragment maturation can leave post-replicative single-stranded gaps behind a fork that keeps moving (37). Leading-strand synthesis persists to higher drug concentrations, but once Pol ε is inhibited, CMG unwinding becomes uncoupled from DNA synthesis (38). The exposed single-stranded DNA activates ATR-Chk1 (39) and promotes RAD51-dependent fork reversal (40). Nor is the fork’s standard rescue route available: leading-strand stalling is normally tolerated by replicase switching to Pol δ (41), yet Pol δ is more sensitive to aphidicolin than Pol ε. No replicative polymerase is spared — all three bind aphidicolin in a conserved pocket (Figure 6), and their IC50 values span roughly one order of magnitude (18). Continued synthesis therefore depends on polymerases that aphidicolin does not inhibit. Pol η and Pol κ cannot sustain processive strand synthesis, but the gaps left by stalled Pol δ are short: the two TLS polymerases exchange with the Pol δ holoenzyme to complete common fragile site synthesis in vitro (42). Fragile sites nevertheless remain unstable in aphidicolin-treated cells, and the structural variants that arise form predominantly after mitotic entry (43). Even so, the stress that aphidicolin produces is dose-dependent and reversible on drug removal. The drug traps polymerases in the pre-catalytic state visualized in our structures (Figure 1-3) rather than damaging DNA or the replisome, which is what makes it a controlled way to perturb DNA replication.

### Toward next-generation B-family polymerase inhibitors

Our structures define the aphidicolin pharmacophore in the assembled holoenzyme. In Pol δ the inhibitor is held by L606, Y607, and D757, with the scaffold packed against Y607 and the D-ring O17 contacting D757 and the L606 backbone. Pol ε interacts through Y631, N813, Y816, and G817. Although hydrogen bond differ in the two reconstructions, equivalent residues occupy every contact position in both enzymes and in Pol α (Figure 6a), the only substitution being L606 in Pol δ for M630 in Pol ε. Aphidicolin therefore inhibits all three replicative polymerases through a conserved B-family architecture, and no scaffold modification has reported to produce a selective derivative. Selectivity cannot come from the residues that touch the inhibitor.

The structures also reframe the existing structure-activity data. Of seventeen A-ring and D-ring derivatives tested against Pol α, none exceeded aphidicolin, A-ring alterations cost more activity than D-ring alterations, and the C-18 function proved critical (44). A-ring C-3 behaves differently. 3-Deoxyaphidicolin inhibits Pol α with a Ki comparable to that of aphidicolin in the same system (45), and a recently isolated congener carrying an ether bridge between C-3 and C-6 retains cytotoxic activity (46). The C-3 oxygen is thus presumably not required for inhibition, and the position tolerates substantial elaboration. Our structures explain this tolerance. The D-ring hydroxyl O17 and A-ring O18 make the anchoring polar contacts in at least two of the three replicative polymerases and can accept little change. O3, however, faces the extruded templating base and makes a single, enzyme-specific contact, to the Pol ε main chain, and is therefore dispensable (Figure 2-3, and 6). Modelling all four bases at the flipped T0 position gives no clashes in either holoenzyme (Figure S3), consistent with competition against all four dNTPs (16, 36) and with the base-independent binding measured for Pol ε (16). Elaboration at C-3 would therefore not be restricted to particular template sequences. In addition, O16, which makes no polar contact in all polymerases, offers a second tolerant vector.

Selectivity must then come from outside the pocket, and the template grip is where the enzymes differ. This is a composite protein-DNA surface rather than a catalytic one, and each polymerase builds it differently. Pol δ holds two template bases beyond T0 through R549, G550, Q551, and Q552 in both the inhibited and nucleotide-bound states (7), whereas Pol ε makes no ordered template contacts beyond T0 in the open state and anchors the template only when the fingers close. Two opposite design objectives follow. A substituent reaching the Pol δ grip would gain contacts that Pol ε cannot offer, a route to a δ-selective compound. Pol α, which likewise lacks an ordered upstream template, is excluded by the same logic. For Pol ε the opportunity is the inverse. Its resistance comes from having nothing to hold the template once the fingers open, so a substituent that contacts the upstream template would supply the clamp the enzyme lacks, and would raise potency where Pol δ, which already has one, would gain little. This mode of inhibition is already druggable. The non-nucleoside inhibitor PNU-183792 binds the active site of the elongating herpesvirus polymerase, displaces the template strand, holds the fingers open and blocks nucleotide binding, with its potency turning on a single contact residue (47). Aphidicolin works in the same mode but lacks this tunability, and the structures presented here show where it could be added, in the grip-directed contacts that derivatives could exploit while keeping the aphidicolin mechanism.

## Data Availability

Consensus cryo-EM maps and atomic models of human Pol δ and Pol ε holoenzymes bound to aphidicolin have been deposited in the EMDB and PDB under accession codes EMD-83585 & PDB-45WK, and EMD-83584 & PDB-45WJ, respectively. The composite cryo-EM map and atomic model of the human Pol δ holoenzyme bound to aphidicolin have been deposited under accession codes EMD-83693 and PDB-46CQ, respectively. The locally refined cryo-EM map of PCNA from the Pol δ holoenzyme has been deposited under accession code EMD-83671.

## Acknowledgements

This research was supported by King Abdullah University of Science and Technology (KAUST) through core funding (to S.M.H. and A.D.B.). We thank L. Zhao, A. Al-Alamoudi, A. Genovese and R. Sougrat for their assistance at the Imaging and Characterization Core Lab at KAUST, and N. Kathiresan and the KAUST IBEX team for access to high-performance computing resources and support with data processing.

## Author contributions

A.U.D. and A.D.B. conceived the study. A.D.B. supervised the research. M.T. prepared the proteins. A.U.D., with the help of S.A. and L.Z., acquired and processed the cryo-EM data and built the atomic models. V.-S.R. and M.T., under the supervision of S.M.H., performed and analysed the inhibition assays. A.U.D. and A.D.B. wrote the manuscript with input from all other authors.

