## Supplementary information for "Structural basis for aphidicolin inhibition of human processive DNA synthesis"

### **SI Guide**

\* These authors contributed equally

#### **Contents**

• Figures S1–4
• Tables S1–2

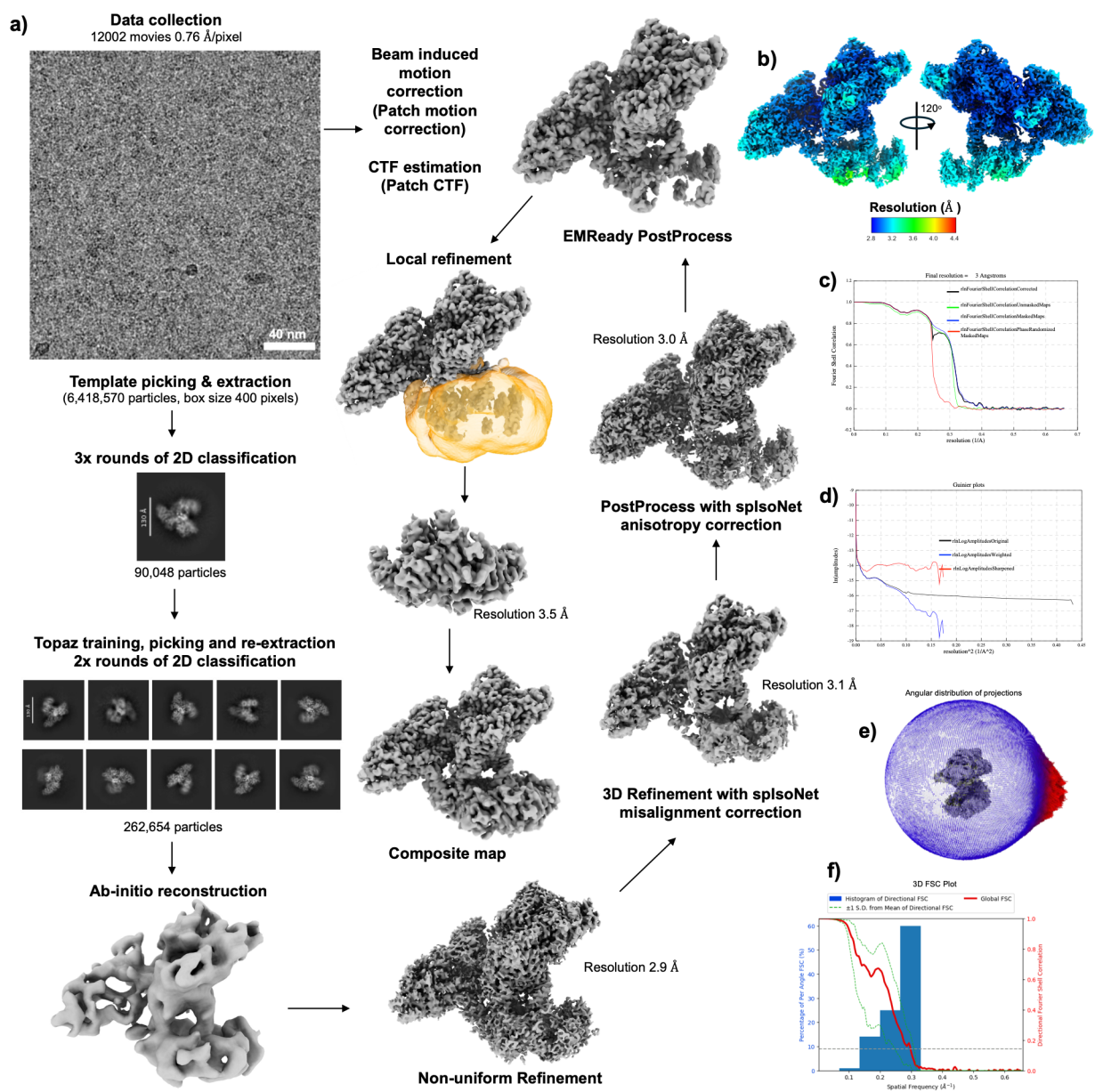

**Figure S1: Cryo-EM data collection and processing of the aphidicolin-bound Pol $\delta$ holoenzyme.** **(a)** Workflow schematics for cryo-EM data collection and processing. **(b)** cryo-EM map colored by local resolution. **(c)** Fourier shell correlation curves for the masked, unmasked, tight-mask, and loose-mask refinements, with resolution estimated using the 0.143 criterion. **(d)** Guinier plot. **(e)** Angular distribution of projections. **(f)** Map isotropy analysis computed by 3DFSC.

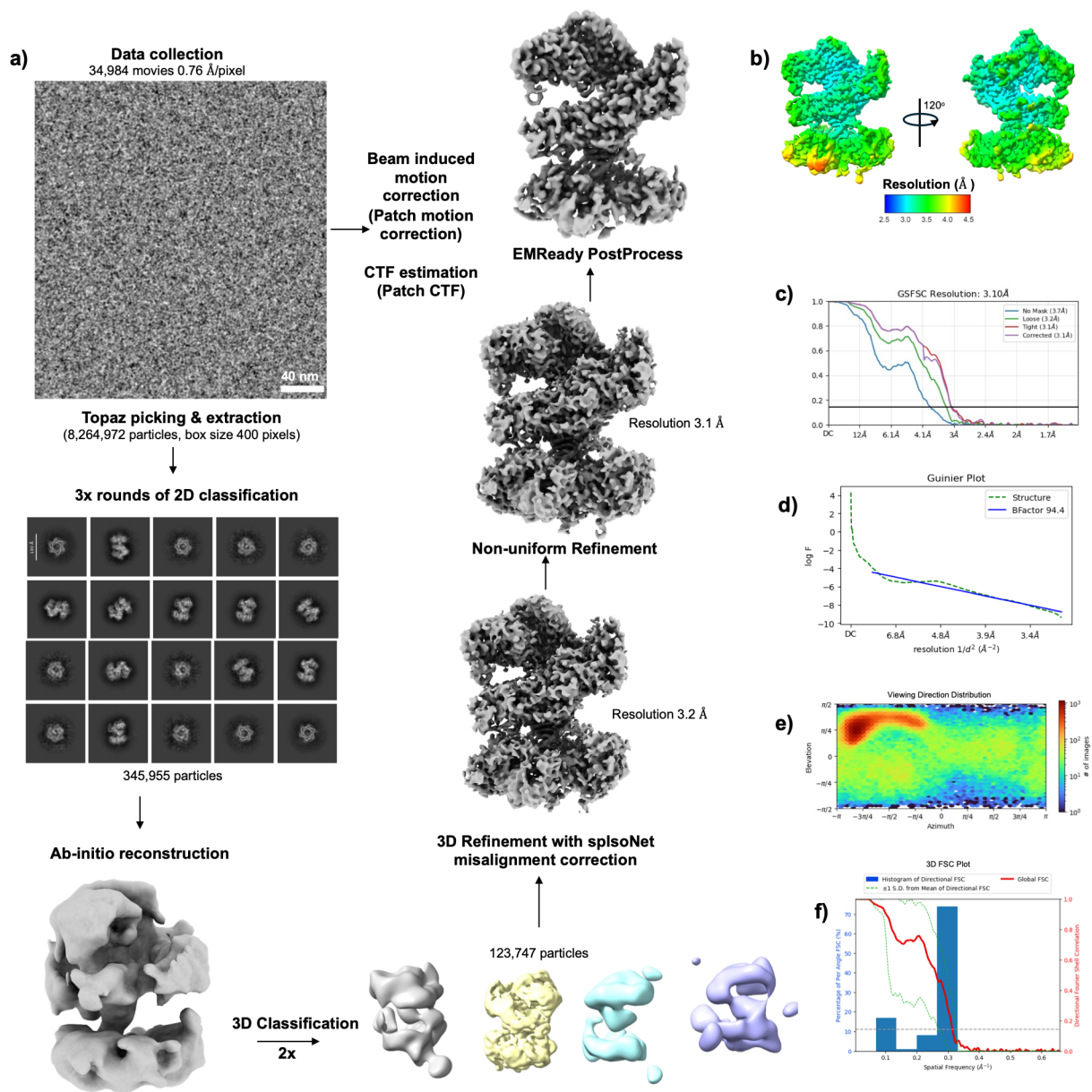

**Figure S2: Cryo-EM data collection and processing of the aphidicolin-bound Pol $\epsilon$ holoenzyme.** (a) Workflow schematics for cryo-EM data collection and processing. (b) cryo-EM map colored by local resolution. (c) Fourier shell correlation curves for the unmasked, tight-mask, loose-mask, and corrected-mask refinements, with resolution estimated using the 0.143 criterion. (d) Guinier plot. (e) Viewing direction distribution. (f) Map isotropy analysis computed by 3DFSC.

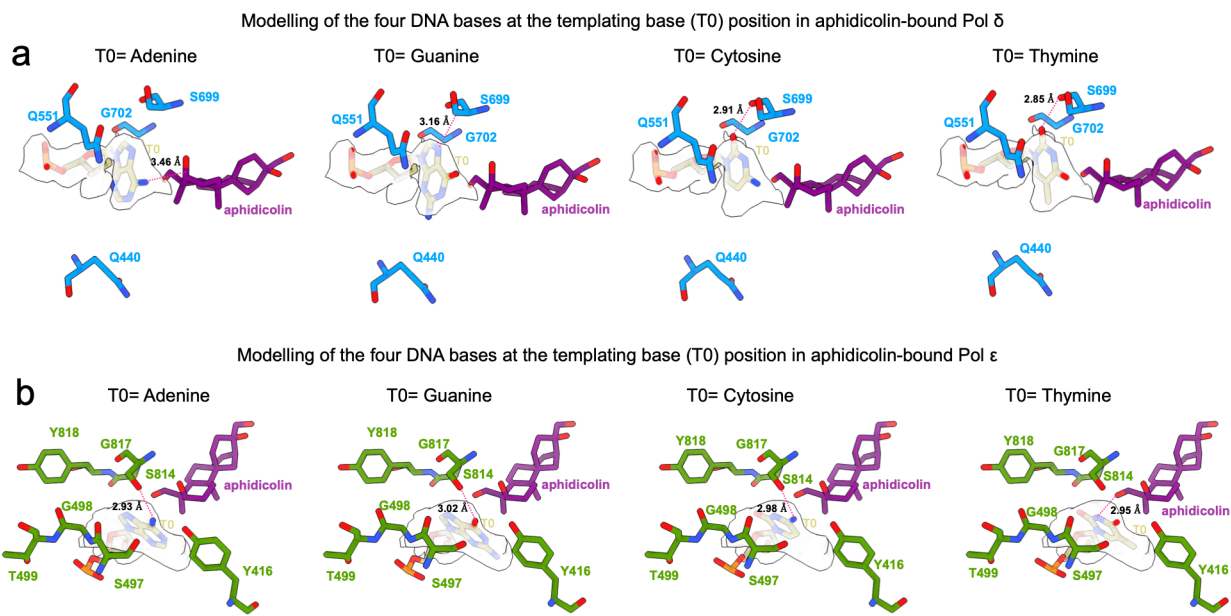

**Figure S3: Modelling of the four DNA bases at the templating base (T0) position in aphidicolin-bound Pol $\delta$ and Pol $\epsilon$.** To test whether aphidicolin inhibition depends on the identity of the templating base, we modelled adenine, guanine, cytosine, and thymine in turn at the flipped-out T0 position in both structures and examined the closest contacts between the modelled base and the surrounding protein and inhibitor. **(a)** Modelled bases at T0 in aphidicolin-bound Pol $\delta$, with closest heavy-atom distances indicated. **(b)** As in a, for aphidicolin-bound Pol $\epsilon$. In both polymerases, no atomic clashes were observed between any modelled base and the protein or inhibitor, suggesting that aphidicolin inhibition does not depend on the identity of the templating base.

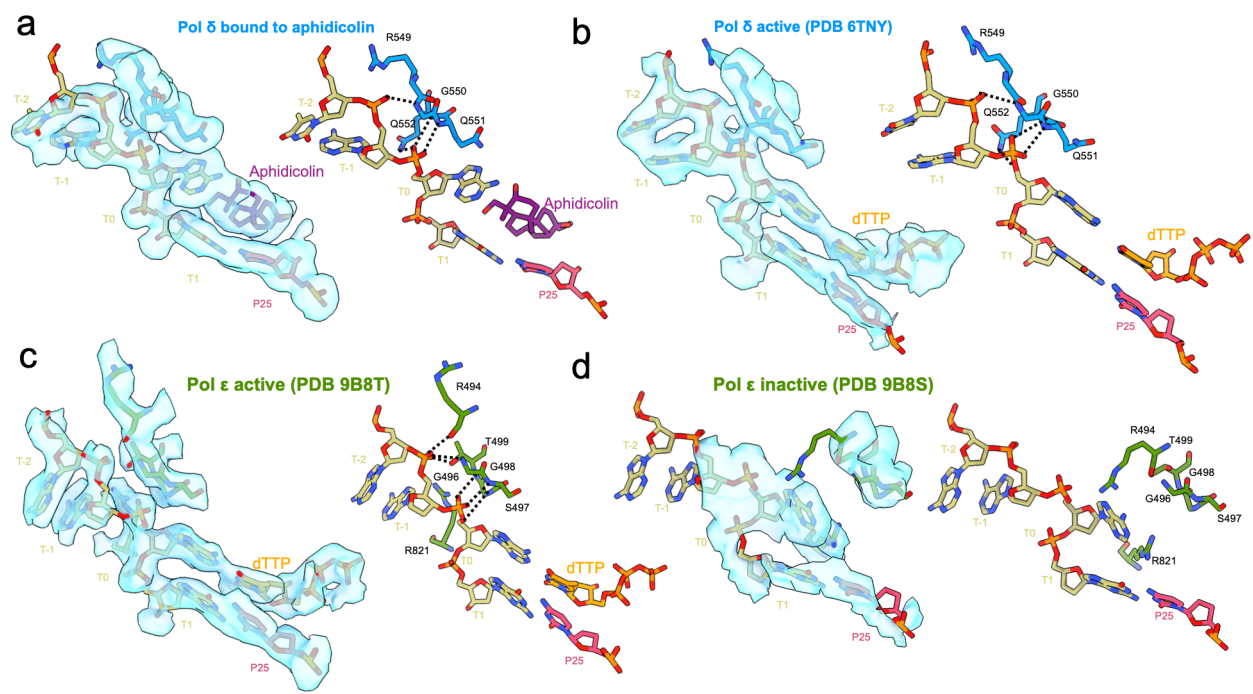

**Figure S4: Comparison of the primer-template junction in aphidicolin-bound and active Pol $\delta/\epsilon$.** To validate the template-strand contacts observed in the aphidicolin-bound structures, we compared the primer-template junction with the active and inactive reference states of both polymerases. **(a)** In the aphidicolin-bound Pol $\delta$ holoenzyme (this study), cryo-EM density shows aphidicolin bound at the templating (T0) position with the T0 base flipped out, while R549, G550, Q551, and Q552 (blue) contact the phosphate backbone of template bases T-2 and T-1 (dashed lines). **(b)** The dTTP-bound Pol $\delta$ ternary complex (PDB 6TNY) (1) shows the same grip residues (blue) making equivalent contacts with the template strand, indicating that aphidicolin binding does not disrupt the template grip. **(c)** The dTTP-bound Pol $\epsilon$ ternary complex (PDB 9B8T) (2) shows template-strand contacts formed by R494, G496, G498, T499, and S497 (green), with R821 positioned near the junction. **(d)** In the inactive Pol $\epsilon$ structure (PDB 9B8S) (2), residues anchoring template beyond T0 retracted to primer-template junction.

**Table S1 – Cryo-EM data collection, model refinement, and validation statistics**

| Polδ-holoenzyme-aphidicolin |  |  |  |
| --- | --- | --- | --- |
| Data collection and processing |  |  |  |
| Magnification |  | 165,000 |  |
| Voltage (kV) |  | 300 |  |
| Total exposure (e-/Å²) |  | 42.62 |  |
| Defocus (µm) |  | -3.0 to -1.0 |  |
| Pixel size (Å) |  | 0.76 |  |
| Symmetry imposed |  | C1 |  |
| Initial particles images |  | 6,418,570 |  |
| Final particles images |  | 262,654 |  |
| Map resolution (Å) |  | 3.0 |  |
| FSC threshold |  | 0.143 |  |
| Map resolution range (Å) |  | 2.8 – 5.0 |  |
| Refinement |  |  |  |
|  |  | Consensus map<br>(EMD-83585)<br>(PDB 45WK) | Composite map<br>(EMD-83693)<br>(PDB 46CQ) |
| Initial model used (PDB code) |  | 6TNY | 6TNY |
| Model resolution (Å) |  | 3.0 | 3.1 |
| FSC threshold |  | 0.5 | 0.5 |
| Map sharpening B factor (Å²) |  | -85.7106 | - |
| Model composition | Non-hydrogen atoms | 19,718 | 19,720 |
|  | Protein residues | 2,399 | 2,399 |
|  | Ligands | 4 | 4 |
| B factor (Å²) | Protein | 59.98 | 66.53 |
|  | Nucleotide | 131.00 | 91.90 |
|  | Ligand | 43.45 | 45.34 |
| R.m.s deviation | Bond length (Å) | 0.007 | 0.012 |
|  | Bond angle (°) | 1.346 | 1.627 |
| Validation | Molprobity score | 1.40 | 1.56 |
|  | Clash score | 2.28 | 3.15 |
|  | Poor rotamers (%) | 0.53 | 0.73 |
| Ramachandran plot (%) | Favored (%) | 94.25 | 92.94 |
|  | Allowed (%) | 5.75 | 7.06 |
|  | Disallowed (%) | 0.00 | 0.00 |

**Table S2– Cryo-EM data collection, model refinement, and validation statistics**

| **Polε-holoenzyme-aphidicolin**<br>**(EMD-83584)**<br>**(PDB 45WJ)** |  |  |
| --- | --- | --- |
| **Data collection and processing** |  |  |
| Magnification |  | 165,000 |
| Voltage (kV) |  | 300 |
| Total exposure (e-/Å<sup>2</sup>) |  | 43.34 |
| Defocus (μm) |  | -2 to -1 |
| Pixel size (Å) |  | 0.76 |
| Symmetry imposed |  | C1 |
| Initial particles images |  | 8,264,972 |
| Final particles images |  | 123,747 |
| Map resolution (Å) |  | 3.1 |
| FSC threshold |  | 0.143 |
| Map resolution range (Å) |  | 2.5 – 7.1 |
| **Refinement** |  |  |
| Initial model used (PDB code) |  | 9B8S |
| Model resolution (Å) |  | 3.4 |
| FSC threshold |  | 0.5 |
| Map sharpening B factor (Å<sup>2</sup>) |  | 94.4 |
| Model composition | Non-hydrogen atoms | 16,206 |
|  | Protein residues | 1,906 |
|  | Ligands | 3 |
| B factor (Å<sup>2</sup>) | Protein | 110.56 |
|  | Nucleotide | 188.81 |
|  | Ligand | 78.52 |
| R.m.s deviation | Bond length (Å) | 0.010 |
|  | Bond angle (°) | 1.401 |
| Validation | Molprobity score | 1.84 |
|  | Clash score | 5.23 |
|  | Poor rotamers (%) | 1.84 |
| Ramachandran plot (%) | Favored (%) | 94.77 |
|  | Allowed (%) | 5.23 |
|  | Disallowed (%) | 0.00 |
